# Method comparison studies of telomere length measurement using qPCR approaches: a critical appraisal of the literature

**DOI:** 10.1101/2020.09.04.282632

**Authors:** Alyssa R. Lindrose, Lauren W. Y. McLester-Davis, Renee I. Tristano, Leila Kataria, Shahinaz M. Gadalla, Dan T. A. Eisenberg, Simon Verhulst, Stacy Drury

## Abstract

Use of telomere length (TL) as a biomarker for various environmental exposures and diseases has increased in recent years. Various methods have been developed to measure telomere length. PCR-based methods remain wide-spread for population-based studies due to the high-throughput capability. While several studies have evaluated TL measurement methods, the results have been variable. We conducted a literature review of TL measurement cross-method comparison studies that included a PCR-based method published between January 1, 2002 and May 25, 2020. A total of 25 articles were found that matched the inclusion criteria. Papers were reviewed for quality of methodologic reporting of sample and DNA quality, PCR assay characteristics, sample blinding, and analytic approaches to determine final TL. Overall, methodologic reporting was low as assessed by two different reporting guidelines for qPCR-based TL measurement. There was a wide range in the reported correlation between methods (as assessed by Pearson’s r) and few studies utilized the recommended intra-class correlation coefficient (ICC) for assessment of assay repeatability and methodologic comparisons. The sample size for nearly all studies was less than 100, raising concerns about statistical power. Overall, this review found that the current literature on the relation between TL measurement methods is lacking in validity and scientific rigor. In light of these findings, we present reporting guidelines for PCR-based TL measurement methods and results of analyses of the effect of assay repeatability (ICC) on statistical power of cross-sectional and longitudinal studies. Additional cross-laboratory studies with rigorous methodologic and statistical reporting, adequate sample size, and blinding are essential to accurately determine assay repeatability and replicability as well as the relation between TL measurement methods.

## Introduction

Telomeres, the protective nucleic acid and protein cap found at the end of all eukaryotic chromosomes, have captured the attention of scientists, medical and public health professionals, biotechnology companies, and the media over the last two decades. In 1973, Olovnikov proposed his theory of marginotomy, which reasoned that during DNA replication, DNA polymerase would not be able to completely copy the first DNA segment and, to prevent the loss of critical DNA sequences in genes, a noncoding set of DNA nucleotides would be required to act as a buffer protecting the loss of important, gene-encoding, sequences [1]. Subsequently in 1978, Blackburn et al. first reported the actual DNA sequences of telomeres in yeast [2], followed by the first sequencing of the human telomere in 1988 [3]. The sequencing of telomeric DNA paved the way for the development of methods that measured the length of telomeres, beginning with the first report of telomere length measurement using the Southern blot method for mammalian chromosomes in 1988 [3]. Since then, thousands of papers assessing telomere length (TL) in human cells have been published across a myriad of different scientific fields (Fig 1). As a result of the broad scientific interest in both the role of TL in disease processes and the influence of environmental factors on TL dynamics, the number of studies evaluating TL in human cells continues to increase, in part facilitated by the regular development of new methods and modifications of existing assays.

**Fig 1.**
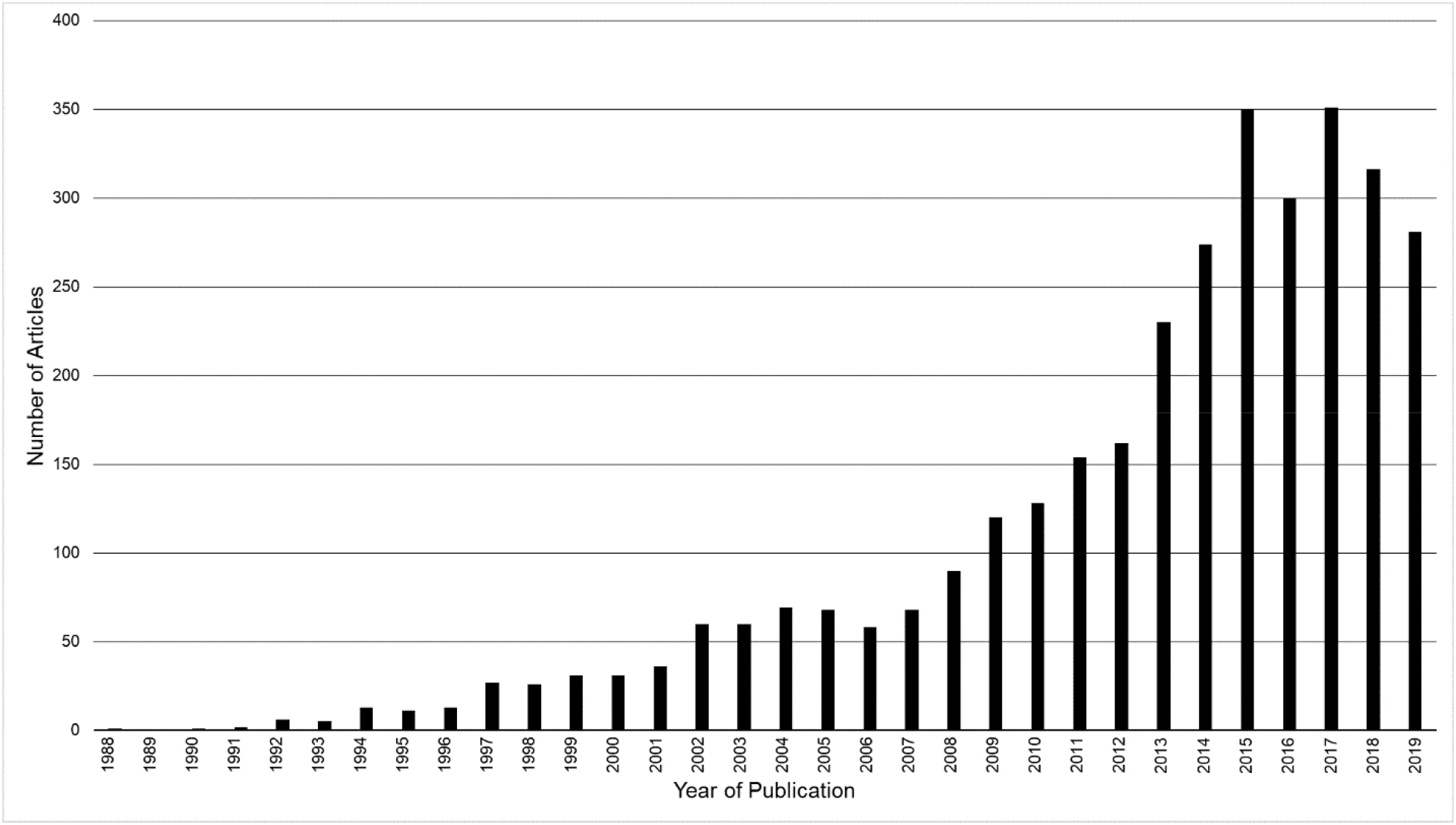
Telomere publications 1988-2019. All publications mentioning telomere length of human DNA from 1988 to 2019 obtained by searching “telomere,” “length,” and “human” in PubMed. Search was completed on May 25, 2020.

Currently, over two dozen assays have been developed to measure TL (Fig 2) [4–34]. These assays are classifiable into four broad categories: hybridization-based, PCR-based, sequence-based, and mixed methods (e.g. hybridization/PCR combination). These assays vary in the information they yield on TL. While most focus on the measurement of the average TL within the sample, assays also measure chromosome-specific TL [5, 16], the complete distribution of TL in a cell population [32], or the shortest TL [25]. The shortest TL has received considerable attention, given evidence from in vitro and preclinical models suggesting that the shortest TL is most predictive of cellular senescence [35, 36]. Several recent reviews have discussed the overall advantages and disadvantages of each method [37–39] focusing on cost, scalability, constraints of starting biological samples (e.g. living cells, amount of DNA, etc) and, to some extent, inter and intra-lab precision as specific challenges facing the field, including the use of coefficient of variation (CV) compared to intraclass correlation coefficients (ICC) [40]. While studies of basic telomere biology continue to explore the complex role that telomeres play in cellular and organismal function, studies testing TL as markers of disease risk or environmental exposure must balance biological relevance, methodologic precision, and experimental practicality, similar to other epigenetic markers such as DNA methylation [41].

**Fig 2.**
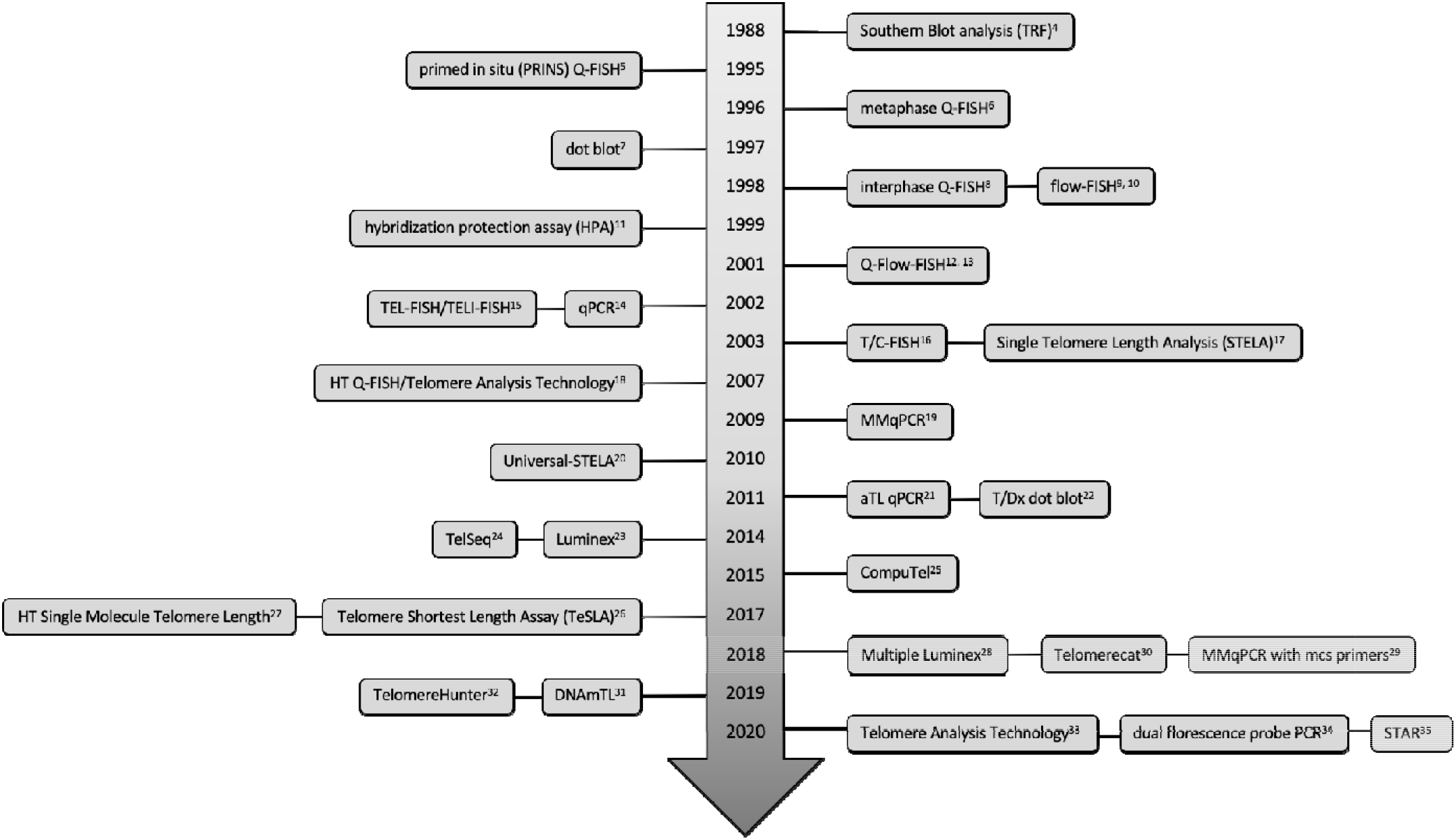
Novel telomere methods developed from 1988 to 2020. Data were obtained by literature search and through references of telomere review papers and method comparison papers.

Over the last decade, debates have arisen over the utility and measurement of TL, particularly with regards to qPCR-based methods. This debate is partially fueled by concerns related to the reproducibility and replicability of TL measurements across studies, methods, and laboratories, and is accentuated by new method development and adaptations of existing protocols without sufficient consensus on the required quality control as more laboratories begin to perform TL assays independently. In response to this debate, several studies have attempted to compare TL measured across different assay methods or laboratories. Some of these method comparison studies examined the direct correlation of TL measurement in the same sample using different assay methods and/or tested the repeatability of TL with the same method (e.g. the amount of within assay variation) [42, 43]. Others tested the relative correlation of the TL measured by different assays with an expected phenotype (e.g. aging, parent-offspring correlation) [44], or examined the relative ability of different assays of TL to predict a specific disease or health outcome [45, 46]. Each of these approaches requires a different analytic strategy and study design and comparison of outcomes is not straightforward. To date, the existing evidence remains insufficient to answer key methodologic questions related to differences in reproducibility and replicability across measurement assays and laboratories, and how/whether these differences affect the ability of TL to serve as a biological indicator of exposure or a predictor of disease or health risk [47]. Beyond these concerns, there remains a lack of consensus as to which, if any, methodology is the “gold standard,” as even the classic Southern blot method is challenged by its inability to capture potentially critical metrics (e.g. full distribution, shortest telomere length, inclusion of the subtelomeric region). To ensure reliability in the widespread utilization of TL as a biomarker of environmental exposure and/or a predictor of a disease, measured by any method that is applicable to population studies, it is critical to systematically test fundamental issues related to assay reproducibility and replicability [48].

As part of a joint National Institute of Aging and National Institute of Environmental Health Science initiative that funded a U24 cooperative award and four separate U01 awards, a Telomere Research Network (TRN) was established in 2019 (trn.tulane.edu). The TRN is coordinating cross-method comparison studies with the long-term goal of developing methodological guidelines and recommendations for telomere research applicable to population-based studies. As a first step towards the goals of this network, we undertook a systematic literature review of published studies that directly compared TL measured using at least one PCR-based method and another approach to determine how these studies might inform the field, with particular attention to assay precision and accuracy of different measurement assays and what research gaps remain.

As defined by the Committee on Reproducibility and Replicability in Science, precision is the closeness of agreement between measured quantities obtained by replicate measurements, while accuracy is the closeness of agreement between a measured quantity and a true value [48]. Reproducibility is defined as precision in measurement under conditions that involve different locations or different measurement procedures, while repeatability is defined as precision in measurements that include the same procedures/locations. Beginning from this perspective, this systematic review evaluated the existing literature related to cross-method comparisons. This review focuses on PCR-based methodologies due to their increasing use in population-based studies, their central role in the debate related to assay precision, and the existence of two reporting guidelines - one created through the TRN (Supplemental Table 1), and a second one created by a separate group in a recently published manuscript [49]. The majority of PCR-based methods are derived from two seminal methodologic papers by Richard Cawthon, the first describing a monoplex based assay (qPCR) and the second describing a multiplex assay (MMqPCR) [13, 18]. Our review focuses specifically on the comprehensiveness of methodologic reporting, correlation between TL measured by different assays, assay repeatability and reproducibility, and overall scientific design of methodological comparisons. Finally, we suggest areas of needed scientific examination and provide some guidance related to study design, necessary sample size, and analytic approach, to address key remaining questions: (1) What is known about the relationship between TL measured using PCR-based methods and other assays? (2) What is known about the reproducibility and repeatability of PCR-based methods and how does this relate to other TL measurement techniques? (3) What are the implications of methodologic precision for sample size and power? (4) What are appropriate guidelines to systematically evaluate the precision of both existing and future TL assays? Addressing these important questions is a requisite step in advancing our understanding of the ability of TL, measured by any approach, to serve as a sentinel of psychosocial and environmental exposures and a predictor of future disease.

## Methods

### Manuscript search

To identify relevant papers that reported on cross method comparisons of any qPCR-based method (qPCR, absolute TL (aTL), and MMqPCR) and another method of TL assessment (PCR-based or otherwise) or the same PCR-based method conducted in separate laboratories, we conducted a critical review beginning with a literature search (Fig 3). The following key terms “telomere,” “telomere length,” and “human” were searched in PubMed and Web of Science. From these initial results, a second search included the keyword “PCR” to identify the initial titles for screening. Search criteria included papers published since January 1, 2002 (the year the first method to measure TL by qPCR was published) through May 10, 2020. The references of selected papers were also reviewed to identify any additional papers. A list of identified papers was presented to the TRN Steering Committee, who also suggested additional papers. Initial review of papers for inclusion was accomplished through evaluation of both the abstract and methods section, as some manuscripts were not directly focused on methodological comparison and instead only reported the cross-method comparison on a subset of samples. Final inclusion in this review met the following baseline requirements:

1. Article published in a peer-reviewed journal (abstracts and pre-prints not included).
2. Article was not presenting the initial development of a new method or a substantial refinement of an existing methodology. This type of study was excluded due to the intrinsic bias in reporting novel method development.
3. Included a direct comparison of TL using the same biological sample measured with two distinct TL assays or the same assay in two or more separate laboratories.
4. At least one of the methods used to measure TL was solely qPCR-based. TeSLA and STELA were not considered due to the additional hybridization component of the assay.

**Fig 3.**
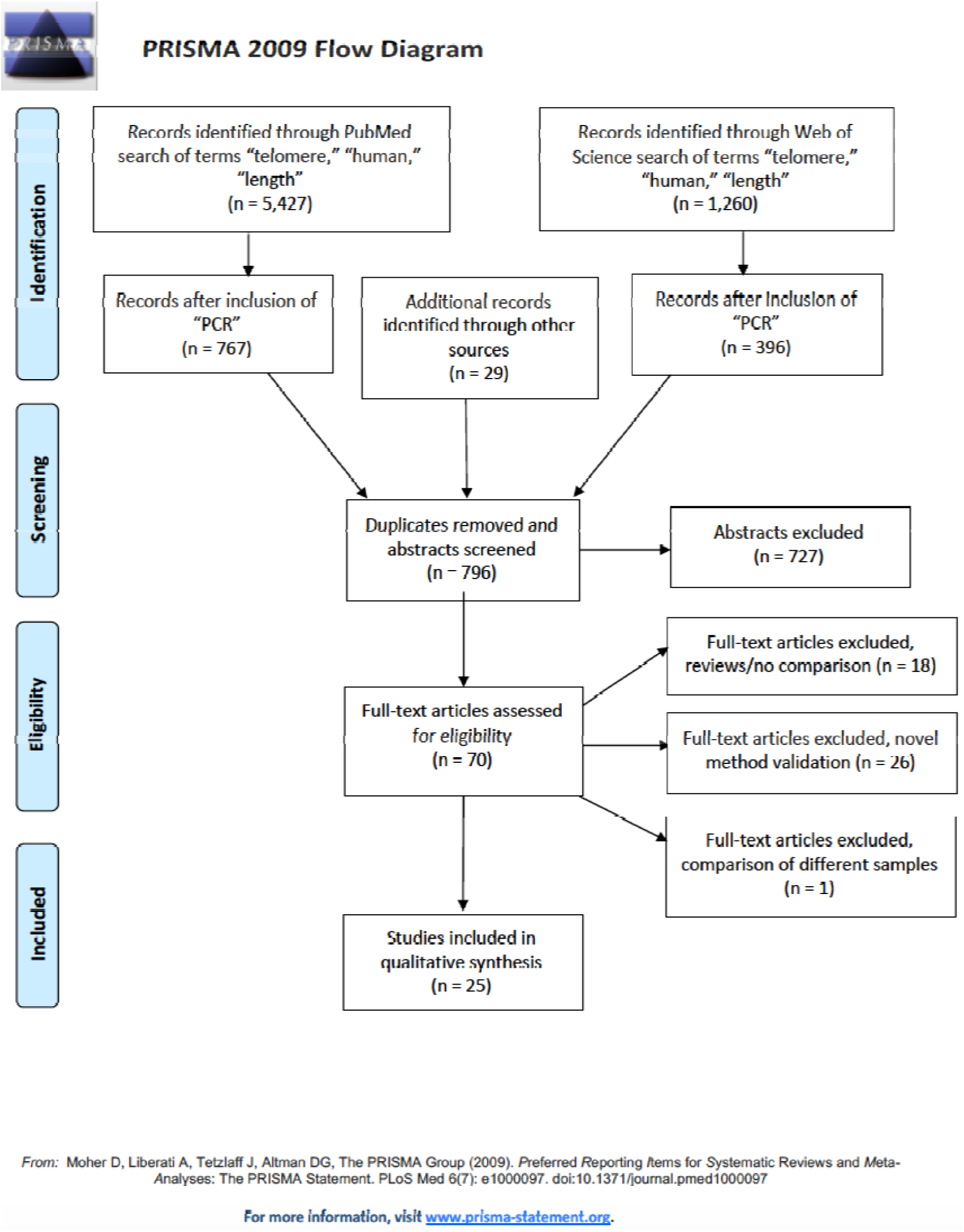
PRISMA flow diagram of literature search and article selection for inclusion. Date range searched was between Jan 1 2002 to May 10 2020. Other sources of article identification included suggestions from participants of the TRN and review of reference lists of selected papers.

### Reporting review

Included papers were evaluated for quality of methodologic reporting using two different indices of reporting guidelines for PCR-based telomere studies. The first was created through consensus of the initial participants in the TRN (Supplemental Table 1). The second was derived from recommendations published by Morinha et al. 2020 (Supplemental Table 2) [49]. We included both guidelines for two reasons. First, there is not empirical data to distinguish between the two guidelines in terms of ensuring rigor and reproducibility for the field. Second, as several of the papers reviewed were authored by participants involved in the creation of the TRN guidelines, the inclusion of both guidelines provided some degree of impartiality. Both guidelines contain overlap with the MIQE guidelines and include characterization of the importance of each recommendation [50]. In terms of specific differences, the Morinha recommendations included several pre-analytic considerations not included in the TRN guidelines (e.g. volume of sample processed, robotic instrumentation vs manual), while storage buffer and the percentage of samples tested for DNA integrity were included in the TRN guidelines but not in the Morinha guidelines. The latter also required greater detail for qPCR validation such as the standard curve and calibration samples, as well as a requirement for the melt curve and Ct of the negative control, which were not included in the TRN guidelines. Both guidelines assess the comprehensiveness of the information describing the PCR assay itself as well as analytic considerations for final TL determination. A grading rubric for each set of guidelines was developed to reduce subjective reviewer interpretation (Supplemental Table 3). A composite assessment for each index was divided into three subcategories for the TRN guidelines and five subcategories for the Morinha guidelines. These broadly encompassed sample collection and processing, DNA quality metrics and storage; PCR assay components and quality control; and data analysis. Two of three reviewers independently assessed each article for fulfillment of reporting guidelines (ARL, LWYM, and SSD). The scores for each individual item were compared and discrepancies resolved by the third reviewer.

Additional characteristics assessed included sample blinding prior to analyses, single lab or multi-lab testing, conversion/transformation of raw TL measurement prior to comparison, and whether the study design evaluated repeatability and/or reproducibility. Lastly, when available, sample size, means and standard deviation of TL measurement are included to assess study power. Although several studies included means and SD of the entire sample, only a subset reported the means and SD of the samples utilized in the method comparison analyses. This review only included method comparisons that involved at least one PCR-based method as currently reporting guidelines are only available for PCR-based methods. As the majority of PCR-based measurements of TL are relative, it was not possible to address accuracy.

Correlation between methods was assessed by using Pearson’s r or r^2^ values where provided. Weighted average correlation coefficients were determined for each type of comparison by converting reported Pearson’s r values (or the square root of reported r^2^ values) to Fisher’s z values, and weighting by sample size. A forest plot was generated from the weighted r average, total sample size for that correlation, and 95% confidence interval (CI) range using DistillerSR Forest Plot Generator from Evidence Partners (https://www.evidencepartners.com/resources/forest-plot-generator/).

### ICC calculation

Given established analytic shortcomings related to the use of the CV as a metric of testing the repeatability of TL, or the correlation of TL measurement between assays, raw data from cross method comparison studies was used to calculate ICCs for comparison between methods where available [40, 51, 52]. ICCs for one study were also calculated using a two-way, single measurement, absolute agreement, random effects model, known as ICC(A,1) and for average measurements ICC(A,k) in McGraw & Wong’s (1996) terminology [53]. The R script used for calculating ICC and associated instructions can be found in the Supplemental Methods.

To provide guidance for future study design, we present several different power analyses outlining the relation between sample size, ICC, and ability to detect group differences. All calculations assumed a realistic (true) standard deviation of 650 base pairs (bp), and N is the combined n of the two groups to be compared and was assumed to be equally distributed among the two groups. Power analysis for cross-sectional comparisons was done using G*Power[54], while power of longitudinal comparisons was estimated through simulations. To examine the impact of variation in ICC on longitudinal TL studies, the statistical power to detect a significant change (paired-t-test) in telomere length of 25 bp/year for sample sizes of 25, 50 and 100 individuals, and an interval of 8 years between baseline and follow-up (i.e. on average 200 bp in total), as a function of measurement repeatability (e.g. reliability) expressed as the ICC. Measurement error was simulated by adding a random number from a normal distribution to the true TL, with the error set at different levels to generate variation in ICC between simulations. Population SD of telomere length was assumed to be 650 bp at both time points and telomere shortening was simulated assuming a Poisson distribution with mean/variance of 25bp/year. This is close to the mean shortening rate typically observed in adults in studies where the age-dependent SD is estimated to be close to 650 bp, and thus the scaling of shortening rate to the overall variance is realistic. Furthermore, power of comparisons using data with another SD can be read from the graphs after rescaling the data to have an SD of 650.

## Results

The initial search revealed 5427 articles and the inclusion of “PCR” as a search term limited the results to 767 articles, whose abstracts and methods were read (Figure 3). An additional 30 articles were identified through assessment of the references of method validation papers and other included cross-method validation studies. A review of these 797 abstracts identified 70 articles for assessment of the full text and supplemental information to determine inclusion in this review. 26 articles were determined to be novel method validation and excluded. 18 articles were excluded as they were either reviews or did not include direct method comparisons. One article was excluded due to the determination that the DNA samples used for cross-method comparison were obtained at different time points. We also included nine papers that, while not specifically designed as a cross method comparison, included sufficient details comparing TL measurement using different assays. This resulted in a total of 25 articles included in this review (Table 1).

**Table 1.**
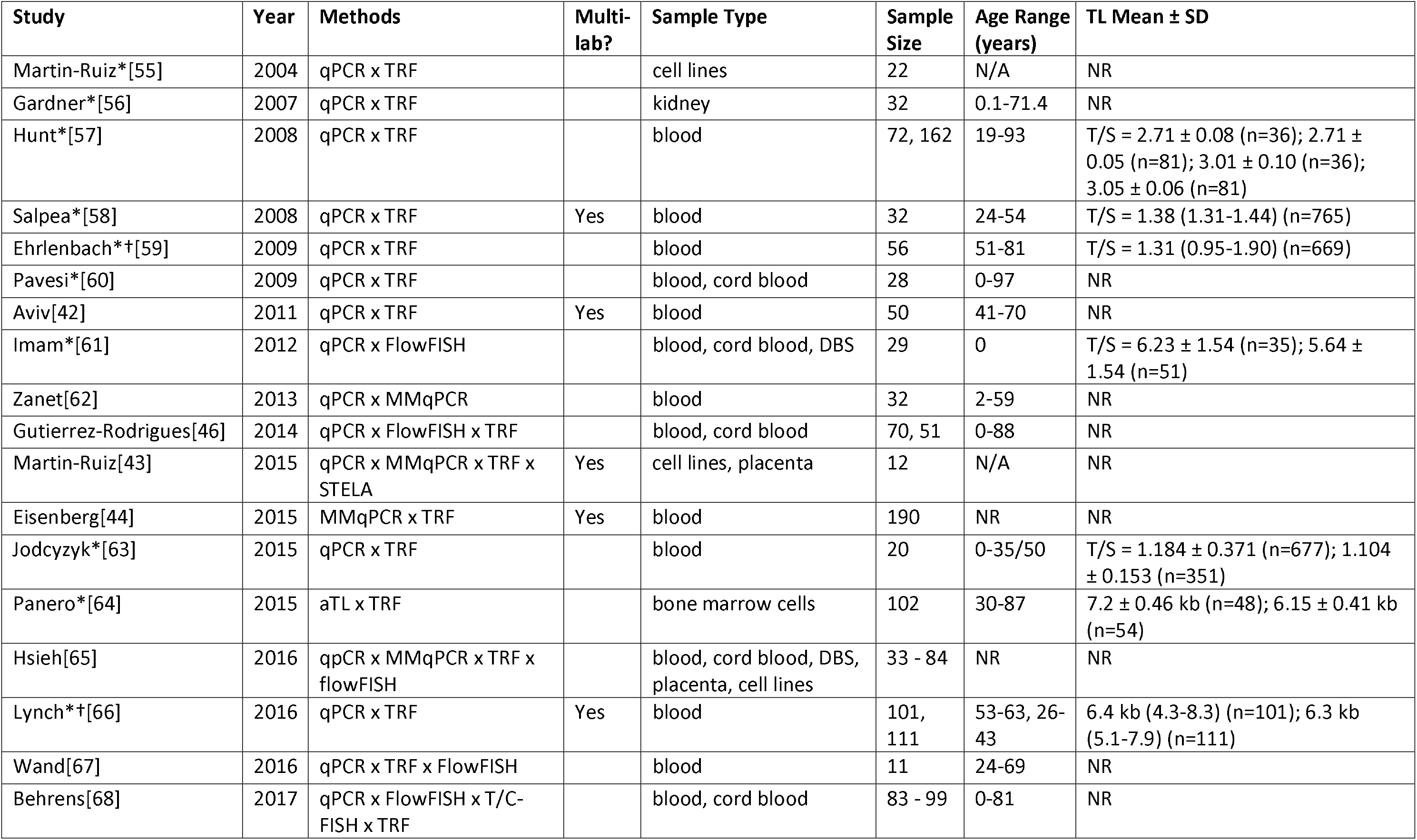
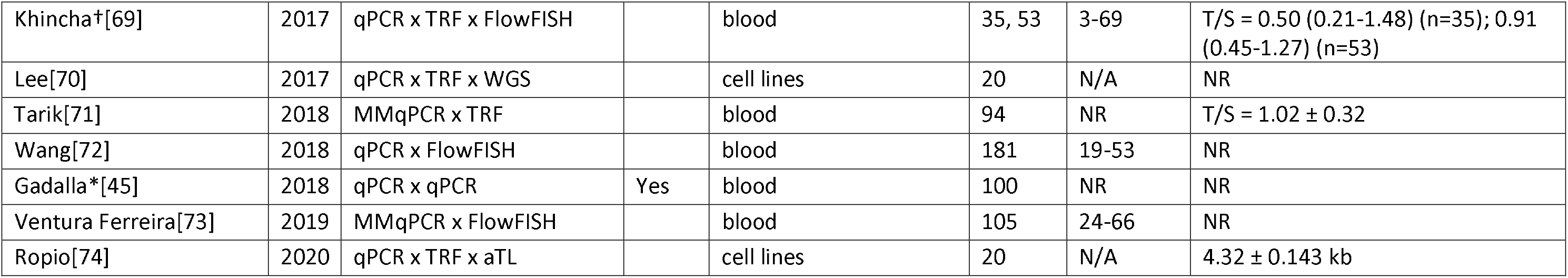
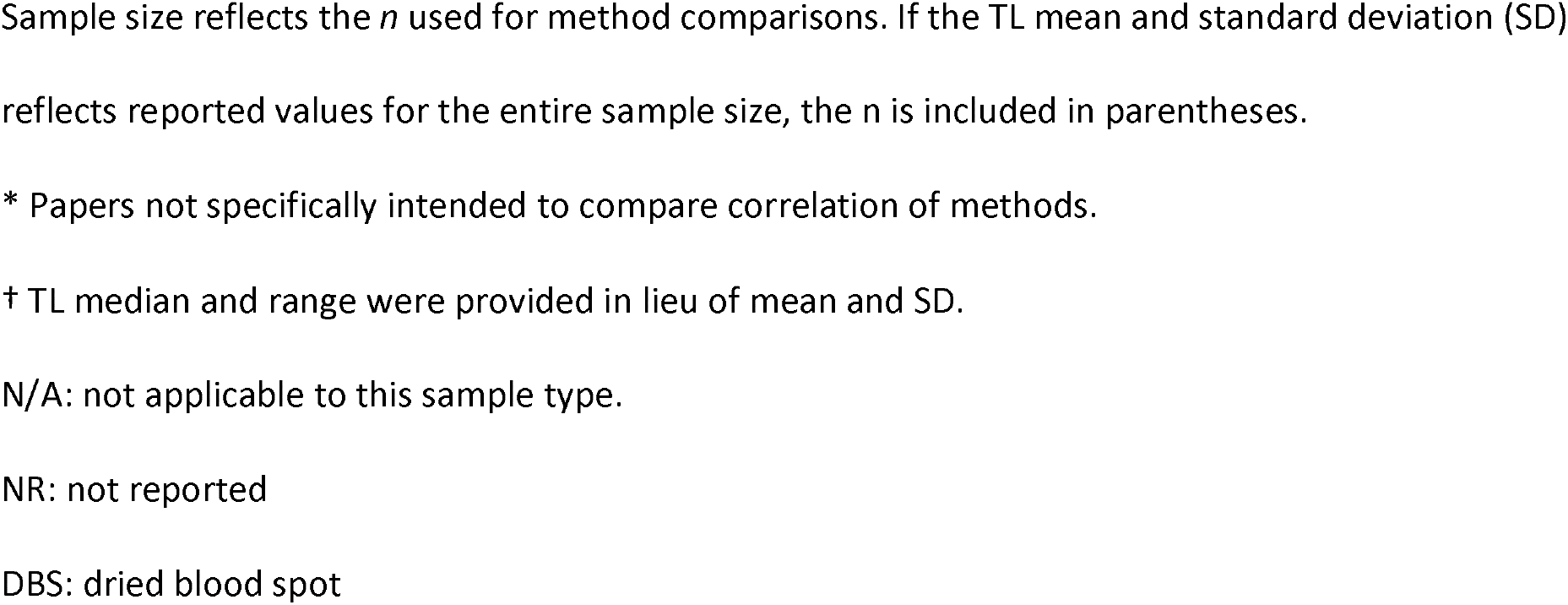
Characteristics of included papers.

### Paper Characteristics

The most common methods comparison among the 25 papers evaluated in this review was monoplex quantitative PCR (qPCR) and the telomere restriction fragment (TRF) method by Southern blot (n=17). Four studies compared multiplex qPCR (MMqPCR) with TRF. Seven studies compared qPCR with the flow-FISH method, and two studies compared MMqPCR with flow-FISH. Three studies involved comparisons of MMqPCR and qPCR, and two studies involved direct comparisons of qPCR methods or separate laboratories. Other methods examined included STELA (compared with qPCR, MMqPCR, and TRF in one study) [43], and absolute TL (two studies, one comparing aTL to TRF and one comparing to aTL to qPCR) [64, 75]. T/C-FISH was also examined in one study as it related to qPCR [68]. Additionally, one paper compared the correlation of several whole genome sequencing (WGS) platforms to qPCR-based measurement[70]. Note some studies compared more than two methods [43, 46, 65, 67–70, 72, 75].

Whole blood was the most common sample type used (n=19), but cord blood (n=5), peripheral blood mononuclear cells (PBMCs) (n=4), and cell lines (n=5) were also utilized as well as a range of other sample types. Several studies reported on more than one sample type.

The reported sample size for the cross-method comparisons ranged from 12 to 181 and only 7 papers had a total sample size greater than 100. Five studies reported the means and standard deviations and two reported the median and range of TL measurements for the study. Two studies provided the raw values of the PCR-based TL measurements.

### Overall quality of reporting of PCR assay methodology

Of the 25 studies included in this systematic review, the average completion score across both reporting guidelines was 51%, with an average of 52% for the TRN guidelines and 50% for the Morinha guidelines (Table 2). Overall, papers included between 26-75% and 29-78% of the recommended reporting metrics for the TRN and Morinha guidelines, respectively. Some metrics were consistently reported in nearly all papers, including the sample type, single copy gene name, and type of PCR method utilized. However, only about 10% of the included papers reported on sample storage, PCR efficiencies, or the number of samples excluded due to quality concerns with the assay.

**Table 2.** Results of reporting guidelines analysis for TRN and Morinha Reporting Guidelines.

|  | Category | Average Score |
| --- | --- | --- |
| TRN Reporting Guidelines | Sample | 37% |
|  | qPCR assay | 75% |
|  | Data analysis | 39% |
|  | **Total** | **52%** |
| Morinha Reporting Guidelines | Sample | 46% |
|  | DNA extraction | 32% |
|  | qPCR validation | 14% |
|  | qPCR protocol | 66% |
|  | Data analysis | 78% |
|  | **Total** | **50%** |
|  | **Average of Both** | **51%** |

Average score reflects the average score across all papers for each reporting category. Total scores reflect the average score across all papers from all reporting items.

### DNA processing

For both the TRN guidelines and Morinha guidelines, reporting of sample type, storage, DNA extraction, and DNA quality/integrity was poor, with an average of 37% for the TRN Sample/DNA category, 46% for the Morinha Sample category, and 32% for the DNA category of the Morinha guidelines. Storage conditions for both the biological samples and extracted DNA were poorly reported, with 24% or less of studies providing this information (Table 3). Fewer than half of the studies reported on metrics related to DNA integrity.

**Table 3.** Item-specific reporting results of the TRN Reporting Guidelines.

| Category | Reporting Item | # Papers Reported |
| --- | --- | --- |
| Sample type, storage, DNA extraction and integrity | Sample type | 25 |
|  | Sample storage temperature | 5 |
|  | Sample storage time before extraction | 2 |
|  | Sample storage buffer | 6 |
|  | DNA extraction method | 20 |
|  | DNA storage conditions | 4 |
|  | DNA freeze-thaw cycles | 4 |
|  | Method of documenting DNA quality/integrity | 8 |
|  | % of samples tested for DNA quality/integrity | 6 |
| qPCR assay | State type of PCR method | 25 |
|  | PCR machine | 21 |
|  | Source of master mix & reagents | 22 |
|  | Final reaction volume | 19 |
|  | Telomere primer sequences | 21 |
|  | Telomere primer concentration | 21 |
|  | Single copy gene name | 25 |
|  | Single gene primer sequence | 22 |
|  | Singe gene primer concentration | 20 |
|  | Full PCR program | 18 |
|  | PCR efficiency of both primers | 3 |
|  | Source of control samples | 14 |
|  | Concentration of DNA standard | 13 |
| Data analysis | Mean and standard deviation/median and range of TL | 13 |
|  | Number of sample replicates | 24 |
|  | Level of independence of replicates | 14 |
|  | Analytic method to determine final TL | 23 |
|  | Method of accounting for variation between replicates | 11 |
|  | Method of accounting for well position effects | 5 |
|  | Method of accounting for between-plate effects | 12 |
|  | % of samples repeated as a result of failed QC | 3 |
|  | % of samples excluded from analysis | 3 |
|  | Acceptable range of PCR efficiency for primers | 6 |
|  | ICCs of sample/study groups | 2 |
|  | T/S ratio transformed to Z score prior to analysis | 2 |

### PCR assay

Reporting on PCR assay conditions and quality control varied. While many metrics of the PCR assay were well-reported, only 18 of 25 studies reported the full cycling conditions. The lowest reporting metric related to PCR was experimental efficiency, with only 12% reporting actual PCR efficiencies. Additionally, just over half (56%) of studies reported the source of their control samples.

### Analytic approaches

Several key reporting gaps were noted in relation to assay quality control and analytic approaches to determining final TL. As with any biologic assay, there is the potential that a specific sample will fail quality control metrics and need to be repeated. Only six papers reported on the number of samples repeated and/or the number of samples that failed quality control. While all but one study reported the numbers of sample replicates, surprisingly, only 14 studies reported on the level of independence of sample replicates (e.g. replicates run on the same plate or on different plates/different times) and only half of the studies reported the means and standard deviations (or median and range) of the T/S ratio.

### Cross-laboratory studies

Only six studies compared analyses across more than one laboratory (Table 1) [42–45]. Of these studies, three described how samples were blinded before analyses. Further, of these cross-laboratory studies, only three studies included the same assay performed in different laboratories [43, 45, 66].

### Reproducibility

Reproducibility, a critical criterion for biologic assays, refers to the relation between measurements using the same assay in different locations or the comparison of values generated using different measurement procedures. This systematic review attempted to assess the relative reproducibility of PCR-based measure of TL in different laboratories as well as the reproducibility precision, e.g. the closeness of two or more measurements, in TL measurement using different methods.

### Relative reproducibility

The current literature does not provide sufficient data to address the relative reproducibility, as, to date, only three studies have tested this directly by performing the same assay in different laboratories or settings. In one study that blinded comparison samples before they were sent to the external laboratories, the median CV across laboratories for qPCR was 18.3%, while the median CV for STELA/TRF based TL measurement was 9.2% [43]. However given the dependence of CVs on the y-intercept, the interpretation of these CVs remains challenging [40]. In the second study, where samples were not blinded before being assayed, the reported within-lab CVs for replicate qPCR measurements were 2.5% and 8.6% [45]. As the laboratories involved utilized different PCR primers, and slightly different methods, it was not possible to directly compare cross-laboratory reproducibility. The third study found inter-assay CVs of 12.0 and 1.2% in two participating labs performing qPCR, but an additional laboratory’s results were excluded from analysis due to an extremely high CV of 27%. Correlation between each laboratory and TRF results were calculated, but no correlation results were provided for the two qPCR assays, and ICC estimates were not reported.

### Reproducibility precision

Reproducibility precision, e.g. the closeness of two or more measurements using different techniques, was addressed to some extent in 19 of the 25 studies reviewed. However, only six studies involved assays performed in different laboratories. The correlation of TL measurement with qPCR-based assays to other assays was, for the most part, reported as linear regression and correlation coefficients (Figure 4). Other papers reported Bland-Altman analyses or did not report a measure of correlation at all. One paper reported mean LTL values for both TRF and qPCR, but did not report a measure of correlation [66]. When examining these results, it should be kept in mind that these methodological studies were generally done in laboratories with extensive experience in the focal technique, and as such are unlikely to be representative of the field at large.

**Figure 4.**
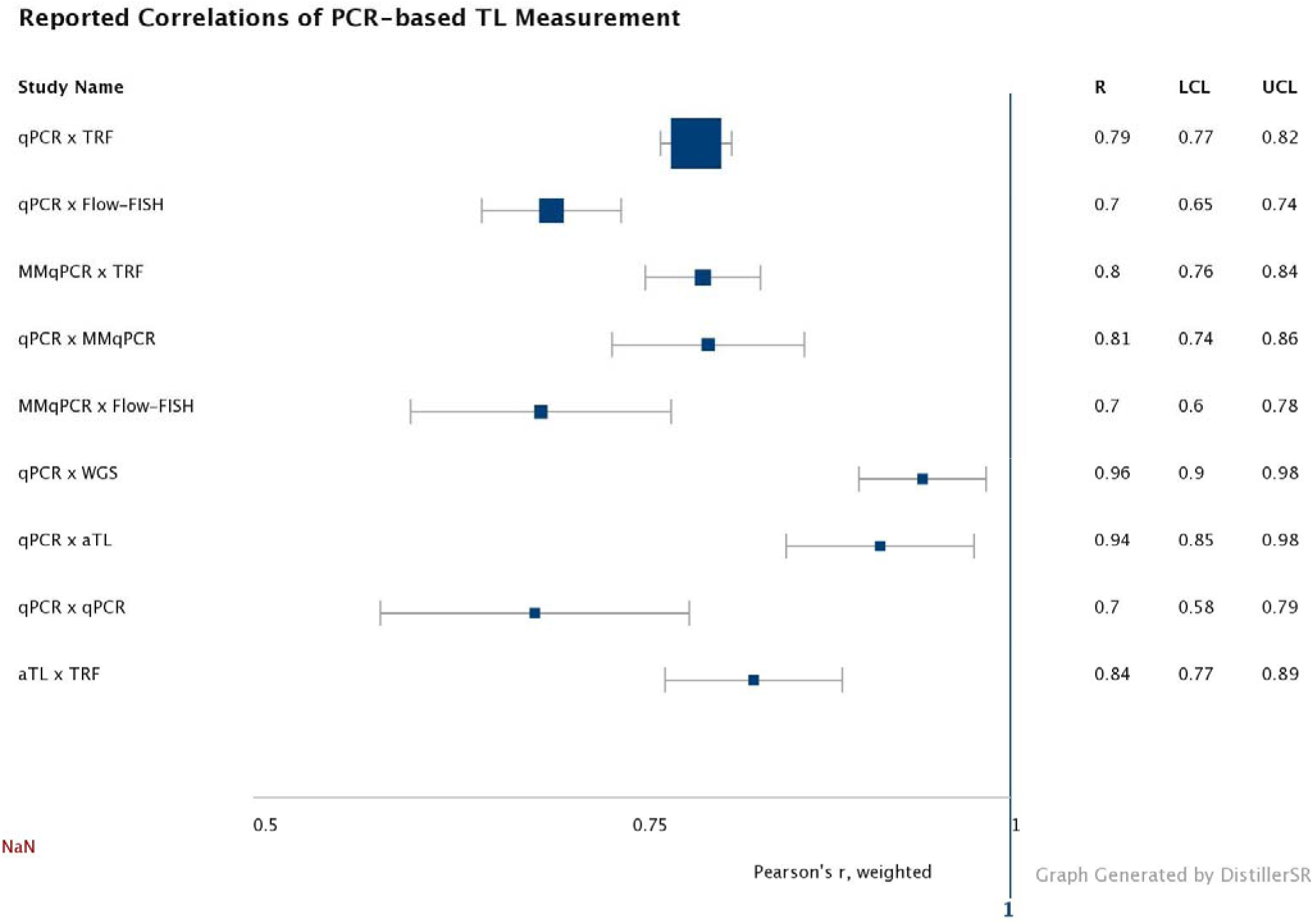
Weighted average correlation (Pearson’s r) and 95% CI for each method compared with qPCR, MMqPCR, or aTL. LCL: lower confidence limit. UCL: upper confidence limit. The point sizes for each comparison are weighted by the number of individual papers with that comparison.

As qPCR and TRF were the most common methods compared, these studies typically reported high correlation, with a weighted correlation coefficient for all studies around 0.75. Correlation of other methods with qPCR or MMqPCR were more variable. No studies have compared MMqPCR to aTL, or whole genome sequencing (WGS). Only one paper each compared qPCR and qPCR (in separate labs), aTL and TRF, qPCR and aTL, or qPCR with WGS. To our knowledge no studies have compared WGS data with aTL, although studies have compared TRF and WGS [23].

In five of the papers in this review, linear regression was used to extrapolate TL into kilobases (kb) from the T/S ratio using TRF values. One paper converted T/S ratio to kb before analysis of the correlation between methods [46]. In two cases, T/S ratio were converted to kb prior to TL comparison utilizing Bland-Altman analysis [68, 73]. In two of the five papers, the conversion of the T/S ratio to bp was based on analyses extrapolated from different data or measured on a different sample type, raising substantial concerns on the true measurement with uncertain implications for the r value [46, 73]. Beyond concerns related to the source of the data utilized for conversion from T/S to bp before comparison across methods, this analytic approach likely to leads to inaccurate reporting [46]. Only two studies utilized the TRN-recommended procedure of transformation to z-score before comparison [43, 66]. When comparing relative TL estimates such as the T/S ratio generated from qPCR, transformation of these values to z-scores will yield more informative results and improve ability to compare results between laboratories or assays [76].

### Repeatability

Repeatability, the precision in measurements that include the same procedure/locations, revealed the greatest variation in both lab and assay specific precision and between methods. In these studies, the number of replicates for a specific DNA sample ranged from 1 replicate (e.g. sample analyzed twice) to five replicates (each sample analyzed six different times). Additionally, only four reviewed studies reported the number of samples that were repeated due to within replicate variance, despite clear acknowledgement in the field that a proportion of all studies will ultimately require repeated assays of telomere length as a result of between-replicate variance. While of limited utility in confirming precision, the intra-assay reported CVs for PCR-based methods (qPCR, MMqPCR) in this review ranged from 2.5 to 12%, and the inter-assay CVs ranged from 3.97 to 15.9%. Inter-assay CVs for TRF ranged from 1.25 to 6.3%, with intra-assay CV reported in only one paper as 1.20%. Inter-assay CVs for flow-FISH were reported as 9.3% and 10.8% in two papers, with only one reporting an intra-assay CV of 9.6%. One paper examining the aTL assay reported its inter-assay CV as 6.7% and intra-assay CV as 2.5%. We emphasize, however, that there are analytic concerns related to the use of CVs for cross laboratory comparisons [40, 52], and directly converting CVs to ICC values is not possible.

Only two studies utilized ICC analyses to examine the repeatability of replicates, reporting ICCs of 0.89 and 0.92 for PCR-based measurement [69, 71]. To expand data on the repeatability of PCR-based and other TL methods, raw data was obtained from authors of a subset of these papers, and ICCs independently calculated. Calculated/reported ICC for TRF methods ranged from 0.92-0.99 in the studies included in this review and are consistent with the ICCs reported in existing studies utilizing the TRF (0.95 to 0.99). However, it is of note that these ICCs were almost entirely the result of TRF measurement in one laboratory. The ICCs for qPCR-based methods in reviewed papers ranged from 0.89-0.92, including the two reported in manuscripts and an additional ICC calculated from raw data (ICC = 0.915, SE = 0.023, 95% confidence interval: [0.860, 0. 946], P<0.001; reported CV 6.5%) [40, 77]. ICCs for MMqPCR (triplicates on the same plate) from one study were run separately based on year of analyses. In one set (n=873) run across different PCR plates in initial and duplicate runs, the ICCs were ICC(A,1) = 0.82 (95% CI 0.79-0.84) and ICC(A,k)=0.90 (95% CI 0.88-0.91). Because these samples were re-run due to initially high intra-assay CVs, this is possibly an under-estimate of the true ICC value. For these same samples, TRF ICCs were calculated from duplicate gels on a subset (n=159) and the inter-gel ICC = 0.96 (95% CI 0.94-0.97). However, we note that these TRF analyses were conducted by a trainee which likely decreased repeatability compared to what is typical of experienced technicians. Given the significant variation in methodologic and raw data reporting, and the wide variability in published CVs, it is likely that the majority of existing TL studies not specifically comparing methods would have significantly lower ICCs.

### Determination of effects of ICC variability on sample size and study power

Our systematic review revealed wide variation in TL measurement repeatability. No biologic assay is perfect, and laboratories measuring any biologic substrate vary in their own internal quality control and repeatability. To provide general guidance for investigators, we therefore conducted analyses to evaluate the impact on power and sample size across a range of ICCs.

In Figure 5A, we present the sample size required to test effect sizes of 150, 200, and 300 bp with a t-test with a power of 0.9, as a function of measurement error as expressed in the ICC. To contextualize the differences: 150 bp is the approximate difference found between the sexes, and 300 bp is the approximate difference observed between individuals with and without atherosclerotic cardiovascular disease [78]. As directly converting bp to T/S ratios is not feasible in this analysis and the analyses below, we suggest that investigators using T/S or other relative TL measurements use standard deviation (SD) differences to estimate power. For example. a difference of 150 bp is equal to 150 / 650 = 0.23 SD, which can be converted to a T/S difference when the SD of the T/S measurements is known. Estimates of potential difference can be extracted from existing literature related to their exposure or outcome of interest when considering study design and sample size.

**Fig. 5A.**
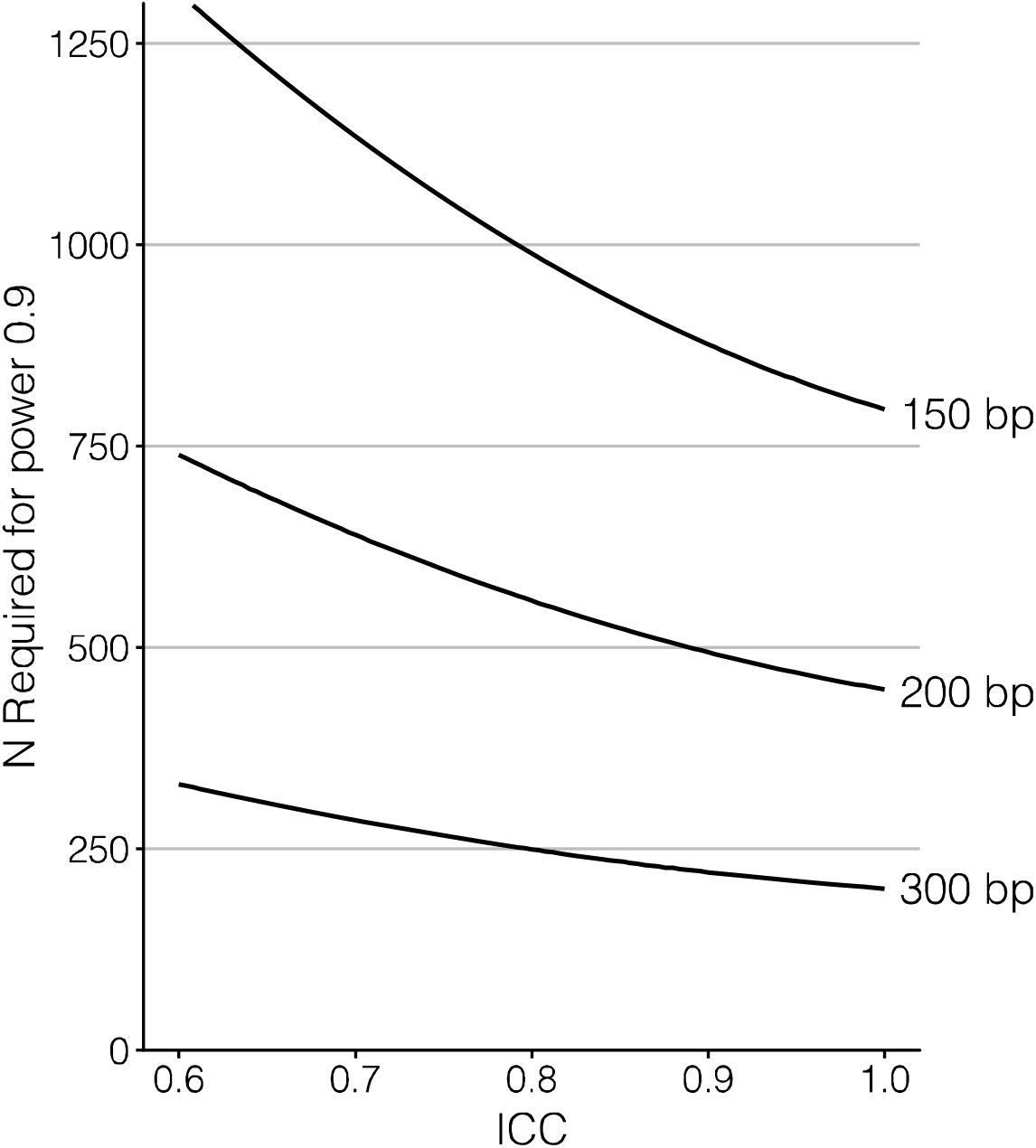
The sample size required to test effect sizes of 150, 200 and 300 bp with a t-test with a power of 0.9, as a function of measurement error as expressed in the ICC. Calculations assumed a realistic (true) standard deviation of 650 bp and power analysis was done using G*Power [54]. N is the combined n of the two groups to be compared and was assumed to be equally distributed among the two groups.

Finally, we present the statistical power of different sample sizes to detect a significant difference in telomere shortening rate of 33% using longitudinal data, as a function of measurement reliability expressed as ICC (Fig. 5B, C). This analysis revealed that even with a high ICC (>0.9), large sample sizes are required to yield sufficient statistical power to detect even substantial differences in telomere shortening rate, in particular when the follow-up period is short. This is due to the rate of telomere shortening being low (here 25 bp/year) compared to the TL variation between individuals (here an SD of 650). The rate of base pair loss in infants and children is likely significantly different and, but as of yet is poorly characterized (but see [79]).

**Fig. 5B, C.**
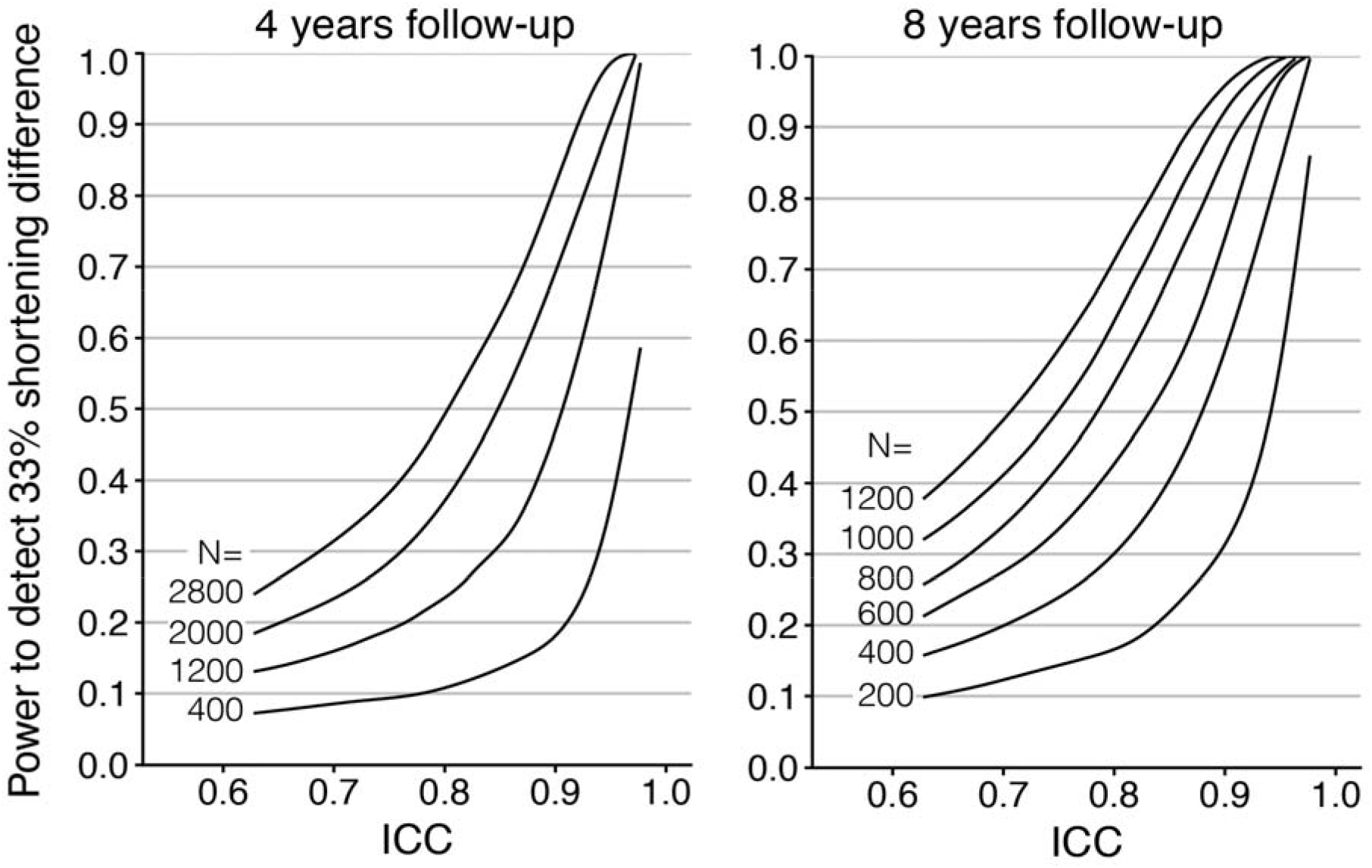
Power to detect a 33% change of telomere shortening rate, up or down, with p<0.05 relative to a baseline shortening rate of 25 bp/year. D. Four-year follow-up period. E. Eight-year follow-up period. Power was calculated for sample sizes as shown (200 – 2800), equally divided over the two levels of telomere shortening rate. Baseline telomere shortening was simulated assuming a Poisson distribution with mean/variance of 25, and population SD of telomere length was maintained at 0.65 kb at both time points.

## Discussion

This systematic review found a total of 25 papers documenting comparison between TL measured using a PCR-based methodology and another TL assay. Until recently, no publication reporting guidelines existed for qPCR-based TL measurement. Our review focused on method comparison studies with the expectation that critical assay parameters and methodologic description would be more detailed and specific. Our review, using two separately developed reporting guidelines, found that, on average, only half of the recommended factors were documented, indicating the need for increased methodologic reporting and wider awareness of reporting recommendations. The lowest reporting was related to information about the validation of PCR-based assays outlined in the Morinha guidelines, with only seven papers including any of the recommended factors. PCR efficiencies, a key reporting requirement in both guidelines and the MIQE guidelines, was absent from the majority of papers with only six mentioning the PCR efficiency parameters and only three documenting the actual PCR efficiencies. Given that all PCR-based methods either assume or specifically calculate the PCR efficiency when determining the T/S ratio, and that, in general, the determination of the T/S ratio assumes similar efficiencies for the single gene and the telomeric primers, the absence of this key metric is concerning. Fewer than half of studies failed to comment on key pre-analytic factors, specifically sample storage time and conditions, freeze-thaw cycles, and evaluation of DNA quality and integrity, all potential sources of assay variability for both PCR and non-PCR-based TL assays that may contribute to current debates in the field about the utility of TL [74, 80, 81]. Lastly, the reporting of the number of samples failing initial quality control, repeated, or unable to be assayed was low. In laboratories routinely performing TL measurement using any assay, a certain percentage of samples for each study will require repeating and regularly a small subset may be unanalyzable for various reasons. While it is possible that these factors were considered and monitored, the lack of reporting for this metric heightens the need for increased attention to the proposed reporting recommendations. Moving forward, the widespread dissemination of these qPCR reporting guidelines to study sections, peer reviewers, and scientists represents an important next step in enhancing the scientific rigor of the field.

At this time, evaluation of the existing literature fails to provide sufficient evidence of the relative or precision reproducibility of different TL assays. Our review identified only six studies that included cross laboratory comparisons and, of these, only three evaluated PCR-based assays performed in more than one lab. As the number of laboratories performing TL studies using PCR and other methods continue to increase, the lack of clear data about cross-laboratory reproducibility and the absence of existing DNA standards or other methods to account for cross-laboratory variation substantially limits the ability to characterize relative reproducibility. In terms of reproducibility across different methods (e.g. PCR and TRF, or PCR and FISH), the current variability in findings, particularly when coupled with limited methodologic reporting, highlights the need for additional rigorous and blinded cross laboratory studies that are adequately powered to accurately determine how TL in a population measured using different assays truly relates. Although 17 studies evaluated the relationship between qPCR and TRF, due to the wide range in reported correlations between TL measurements, the relatively small samples sizes, and the insufficient analytic and assay blinding, there is currently insufficient data to draw firm conclusions on the general correlation between TL measured with different assays. The analytic consequences of using CVs to test the relationship between TL measured using different assays has been discussed previously, as has the issues caused by the use of analytic strategies such as conversion to base pairs instead of z scores, especially when extrapolating from data produced in different laboratories or using different samples [40, 51]. In this review, we utilized existing raw data from 3 included studies to provide preliminary data about precision reproducibility for PCR and TRF studies. The wide range of ICCs calculated from these few studies, particularly for PCR-based methods, and the low reporting of ICCs in the papers included in this review highlights the need to increase attention to this specific reporting requirement. For many of the existing studies, the small sample size and the lack of reporting of the means and standard deviations of TL prevents objective determination of whether any of the current studies were adequately powered. Beyond these concerns, the over-representation of data from specialized laboratories, particularly for TRF, the applicability of much of the existing data to the wider telomere field is uncertain. For aTL and MMqPCR-based TL measurements, the current paucity of published cross method comparisons limits the ability to form an opinion of how TL measured with these assays relates to other methods.

Measurement precision is critical, in particular for longitudinal studies. Methodologies that are low cost, practical, and simple to implement with standard laboratory equipment, especially when they are innovative or high impact, are often rapidly implemented across laboratories with various levels of expertise in the new methodology. Invariably this results in diverse protocols, analyses, and methodologic reporting – consequences that are even more problematic when there is an absence of consensus on best practices [50]. As with many other biologic assays, the development of reporting guidelines for TL measurement has lagged behind the broad implementation of the methods themselves [82–86]. The lack of consolidated guidance about factors, both pre-analytic and within the assay itself, that contribute to measurement error when combined with the wide popularity of PCR-based TL measurement undoubtedly contributed to discrepancies in the existing literature and failed study replications. Similar to the MIQE guidelines, the reporting guidelines presented and tested in this systematic review for PCR-based TL assays are meant as minimal reporting recommendations focused on enhancing the reliability of results, consistency between different laboratories performing the same assay, and increased experimental transparency and accuracy [50]. To assist investigators and reviewers we highlight the overlapping recommendations with the MIQE guidelines, indicate whether a particular requirement is desirable or essential, and provide references that support the selection of the particular reporting requirement. Over the course of the next four years, the TRN expects to develop similar reporting recommendations for other types of TL assays while conducting adequately powered and scientifically rigorous studies to support these reporting guidelines, recognizing that individual recommendations have varying levels of initial empirical support [87].

Despite the strengths of this review, there are several limitations. First, this review only focused on assays applicable to population-based studies in humans. It does not address issues in other species or assays that may have clinical utility but for which the requirements for sample types (e.g. fresh tissue and/or live cells) or the cost/labor/expertise requirements (e.g. TeSLA, STELA) limit utilization in population based studies. A second limitation is that we utilized reporting guidelines for qPCR-based assays only. To date, specific protocol recommendations and reporting guidelines have not been published for other TL assays (e.g. TRF, FISH) although detailed methodologic protocols do exist [88]. Additionally, it is possible that additional articles comparing TL assays may be available in other databases or pre-print servers. However, many of the articles included in this review were not specifically designed solely to compare TL measurement methods and would not be found through standardized database searches. Further, it is unlikely that additional articles would change the general picture emerging from this review. Finally, we note that while this article focused on precision and reproducibility, accuracy of measurement is as important. Precisely inaccurate measures will be of limited use to the scientific field, a factor that becomes more problematic when using relative estimates and not true values as is the case in many TL assays. In the absence of a clear gold standard measurement technique, accuracy is difficult to discern.

## Conclusions

After careful examination of the existing literature, it is apparent that rigorous cross laboratory and methodological studies must be an immediate priority for the field. To assist the field moving forward, we include reporting guidelines for PCR-based TL assays and indicate specific scientific papers that support these recommendations originally developed through consensus of the initial TRN members and consultants. These guidelines do not outline a specific PCR methodology and, at this time, we do not believe there is sufficient data to provide guidance on specific assay approaches or components. Rather, these guidelines are provided to ensure reviewers and readers can adequately assess the methodology and consider the implications of these factors for each study’s findings. The consistency in results across reporting guidelines (TRN, Morinha, MIQE) related to the integrity and quality of both the initial biological sample and the DNA itself support the critical nature of this reporting metric. In terms of assay reporting, increased attention of investigators and reviewers to ensuring complete reporting of assay reagents and PCR efficiencies is also expected to enhance the rigor of the field. TRN investigators are currently testing the impact of different pre-analytic factors, DNA integrity, and PCR conditions to provide extant evidence of the importance of these parameters in relation to precision and reproducibility. We recommend that studies be required to report ICCs in lieu of CVs, as well as either the median or mean and standard deviation of TL. We also provide specific guidance related to sample size and power that is contingent upon the ICC given the substantial impact of differences in assay precision on the ability to determine true relationships and with the expectation that this will be of use for investigators as they embark on new research studies. It is important to balance assay cost, in terms of both time and reagents, with the needed sample size and statistical power. Moving forward, investigators should carefully consider study design from this perspective, recognizing that there is currently no “ideal” approach. Telomere research offers significant potential across a diverse range of scientific fields with potential mechanistic insight into overlapping biological pathways contributing to many of the leading causes of morbidity and mortality. Ensuring the highest scientific rigor and precision, through accurate methodological reporting and rigorous testing of the factors that contribute to assay variability, are requisite steps to ensuring that potential is achieved.

## Acknowledgements

The authors would like to acknowledge the TRN Advisory Committee (Daniel Belsky, Eileen Crimmins, Sonja Entringer, Daniel Nettle, Jerry Shay, Daniel Nettle), U01 grantee P.I.s (Abraham Aviv, Jue Lin, Idan Shalev, Yun-Ling Zheng), and NIA/NIEHS partners (Max Guo, Michelle Heacock, Lisbeth Nielsen) for their input in creating the TRN qPCR Reporting Guidelines.

## References

1. Olonikov, A., A theory of marginotomy. J Theor Biol, 1971. 41: p.181–190.

2. Blackburn, E. and J. Gall, A tandemly repeated sequence at the termini of the extrachromosomal ribosomal RNA genes in Tetrahymena. Journal of molecular biology, 1978 120(1): p. 33–53.

3. Moyzis, R., et al., A highly conserved repetitive DNA sequence,(TTAGGG) n, present at the telomeres of human chromosomes. Proceedings of the National Academy of Sciences, 1988 85(18): p. 6622–6626.

4. Therkelsen, A., et al., Staining of human telomeres with primed in situ labeling (PRINS). Cytogenet Cell Genet, 1995 68(1-2): p. 115–8.

5. Lansdorp, P., et al., Heterogeneity in telomere length of human chromosomes. Hum Mol Genet, 1996 5(5): p. 685–91.

6. Bryant, J., et al., Measurement of telomeric DNA content in human tissues. Biotechniques, 1997 23(3): p. 476–8, 480, 482, passim.

7. de Pauw, E., et al., Assessment of telomere length in hematopoietic interphase cells using in situ hybridization and digital fluorescence microscopy. Cytometry, 1998 32(3): p. 163–9.

8. Hultdin, M., et al., Telomere analysis by fluorescence in situ hybridization and flow cytometry. Nucleic Acids Res, 1998 26(16): p. 3651–6.

9. Narath, R., et al., Automatic telomere length measurements in interphase nuclei by IQ FISH. Cytometry Part A: the Journal of the International Society for Analytical Cytology, 2005 68(2): p. 113–120.

10. Nakamura, Y., et al., Simple, rapid, quantitative, and sensitive detection of telomere repeats in cell lysate by a hybridization protection assay. Clin Chem, 1999 45(10): p. 1718–24.

11. Law, H. and Y. Lau, Validation and development of quantitative flow cytometry based fluorescence in situ hybridization for intercenter comparison of telomere length measurement. Cytometry,: The Journal of the International Society for Analytical Cytology, 2001 43(2): p. 150–153.

12. Rufer, N., et al., Telomere length dynamics in human lymphocyte subpopulations measured by flow cytometry. Nat Biotechnol, 1998 16(8): p. 743–7.

13. Cawthon, R., Telomere measurement by quatitative PCR. Nucleic Acids Research, 2002 30(10): p. e47–53.

14. Meeker, A. K., et al., Telomere length assessment in human archival tissues: combined telomere fluorescence in situ hybridization and immunostaining. The American Journal of Pathology, 2002 160(4): p. 1259–1268.

15. Perner, S., et al., Quantifying Telomere Lengths of Human Individual Chromosome Arms by Centromere Calibrated Fluorescence in Situ Hybridization and Digital Imaging. The American Journal of Pathology, 2003 163(5): p. 1751–1756.

16. Baird, D., et al., Extensive allelic variation and ultrashort telomeres in senescent human cells. Nat Genet, 2003 33(2): p. 203–7.

17. Canela, A., et al., High throughput telomere length quantification by FISH and its application to human population studies. Proc Natl Acad Sci U S A, 2007 104(13): p. 5300–5.

18. Cawthon, R., Telomere length measurement by a novel monochrome multiplex quantitative PCR method. Nucleic Acids Research, 2009 37: p. e21.

19. Bendix, L., et al., The load of short telomeres, estimated by a new method, Universal STELA, correlates with number of senescent cells. Aging Cell, 2010 9(3): p. 383–97.

20. O’Callaghan, N. and M. Fenech, A quantitative PCR method for measuring absolute telomere length. Biological procedures online, 2011 13: p. 3–3.

21. Kimura, M. and A. Aviv, Measurement of telomere DNA content by dot blot analysis. Nucleic Acids Research, 2011 39(12): p. e84–e84.

22. Kibriya, M., et al., Measurement of telomere length: a new assay using QuantiGene chemistry on a Luminex platform. 2014, AACR.

23. Ding, Z., et al., Estimating telomere length from whole genome sequence data. Nucleic Acids Res, 2014 42(9): p. e75.

24. Nersisyan, L. and A. Arakelyan, Computel: Computation of Mean Telomere Length from Whole Genome Next Generation Sequencing Data. PLOS ONE, 2015 10(4): p. e0125201.

25. Lai, T., et al., A method for measuring the distribution of the shortest telomeres in cells and tissues. Nature communications, 2017 8(1): p. 1–14.

26. McCaffrey, J., et al., High throughput single molecule telomere characterization. Genome research, 2017 27(11): p. 1904–1915.

27. Jasmine, F., et al., A novel pooled sample multiplex luminex assay for high throughput measurement of relative telomere length. American Journal of Human Biology, 2018 30(4): p. e23118.

28. Dahlgren, P., et al., Development of a New Monochrome Multiplex qPCR Method for Relative Telomere Length Measurement in Cancer. Neoplasia, 2018 20(5): p. 425–431.

29. Farmery J. H R., et al., Telomerecat: A ploidy agnostic method for estimating telomere length from whole genome sequencing data. Scientific Reports, 2018 8(1): p. 1300.

30. Lu, A., et al., DNA methylation based estimator of telomere length. Aging, 2019 11(16): p. 5895–5923.

31. Feuerbach, L., et al., TelomereHunter–in silico estimation of telomere content and composition from cancer genomes. BMC bioinformatics, 2019 20(1): p. 1–11.

32. de Pedro, N., et al., Analytical Validation of Telomere Analysis Technology^®^ for the High Throughput Analysis of Multiple Telomere Associated Variables. Biological Procedures Online, 2020 22(1): p. 2.

33. Sethi, I., et al., Dual Labeled Fluorescence Probe Based qPCR Assay to Measure the Telomere Length. Gene 2020: p. 145178.

34. Luo, Y., et al., Massively parallel single molecule telomere length measurement with digital real time PCR. Science Advances, 2020 6(34): p. eabb7944.

35. Hemann, M., et al., The shortest telomere, not average telomere length, is critical for cell viability and chromosome stability. Cell, 2001 107(1): p. 67–77.

36. Xu, Z., et al., The length of the shortest telomere as the major determinant of the onset of replicative senescence. Genetics, 2013 194(4): p. 847–857.

37. Montpetit, A., et al., Telomere length: a review of methods for measurement. Nursing research, 2014 63(4): p. 289.

38. Shay,J. W., and W.E. Wright, Telomeres and telomerase: three decades of progress. Nat Rev Genet, 2019 20(5): p. 299–309.

39. Aubert, G., M. Hills, and P. Lansdorp, Telomere length measurement caveats and a critical assessment of the available technologies and tools. Mutat Res, 2012 730(1-2): p. 59–67.

40. Eisenberg, D., Telomere length measurement validity: the coefficient of variation is invalid and cannot be used to compare quantitative polymerase chain reaction and Southern blot telomere length measurement techniques. International Journal of Epidemiology, 2016 45(4): p. 1295–1298.

41. Ehrlich, M., DNA hypermethylation in disease: mechanisms and clinical relevance. Epigenetics, 2019 14(12): p. 1141–1163.

42. Aviv, A., et al., Impartial comparative analysis of measurement of leukocyte telomere length/DNA content by Southern blots and qPCR. Nucleic Acids Research, 2011 39(20): p. e134.

43. Martin-Ruiz, C., et al., Reproducibility of telomere length assessment: an international collaborative study. International journal of epidemiology, 2015 44(5): p. 1673–1683.

44. Eisenberg, D., C. Kuzawa, and M.G. Hayes, Improving qPCR telomere length assays: Controlling for well position effects increases statistical power. American Journal of Human Biology, 2015: p. n/a–n/a.

45. Gadalla, S., et al., No association between donor telomere length and outcomes after allogeneic unrelated hematopoietic cell transplant in patients with acute leukemia. Bone, marrow transplantation, 2018 53(4): p. 383–391.

46. Gutierrez-Rodrigues, F., et al., Direct Comparison of Flow FISH and qPCR as Diagnostic Tests for Telomere Length Measurement in Humans. PLoS ONE, 2014 9(11): p. e113747.

47. Bodelon, C., S. Savage, and S. Gadalla, Telomeres in molecular epidemiology studies. Prog Mol Biol Transl Sci, 2014 125: p. 113–31.

48. Plant, A., R. Hanisch, and National Institute of Standards and Technology, Reproducibility and Replicability in Science, A Metrology Perspective 2018: A Report to the National Academies of Sciences, Engineering and Medicine Committee on Reproducibility and Replicability in Science.

49. Morinha, F., P. Magalhães, and G. Blanco, Standard guidelines for the publication of telomere qPCR results in evolutionary ecology. Molecular Ecology Resources, 2020 20(3).

50. Bustin, S., et al., The MIQE Guidelines: Minimum Information for Publication of Quantitative Real Time PCR Experiments. Clinical Chemistry, 2009 55(4): p. 611–622.

51. Verhulst, S., et al., Response to: Reliability and validity of telomere length measurements. International journal of epidemiology, 2016 45(4): p. 1298–1301.

52. Nettle, D., et al., Consequences of measurement error in qPCR telomere data: A simulation study. PLoS One, 2019 14(5): p. e0216118.

53. McGraw, K. and S. Wong, “ Forming inferences about some intraclass correlations coefficients”: Correction. 1996.

54. Faul, F., et al., Statistical power analyses using G* Power 3.1: Tests for correlation and regression analyses. Behavior research methods, 2009 41(4): p. 1149–1160.

55. Martin-Ruiz, C., et al., Stochastic variation in telomere shortening rate causes heterogeneity of human fibroblast replicative life span. J Biol Chem, 2004 279(17): p. 17826–33.

56. Gardner, J. P., et al., Telomere dynamics in macaques and humans. J Gerontol A Biol Sci Med Sci, 2007 62(4): p. 367–74.

57. Hunt, S. C., et al., Leukocyte telomeres are longer in African Americans than in whites: the National Heart, Lung, and Blood Institute Family Heart Study and the Bogalusa Heart Study. Aging Cell, 2008 7(4): p. 451–8.

58. Salpea, K. D., et al., The association of telomere length with paternal history of premature myocardial infarction in the European Atherosclerosis Research Study II. J Mol Med (Berl), 2008 86(7): p. 815–24.

59. Ehrlenbach, S., et al., Influences on the reduction of relative telomere length over 10 years in the population based Bruneck Study: introduction of a well controlled high throughput assay. International Journal of Epidemiology, 2009 38(6): p. 1725–1734.

60. Pavesi, E., et al., Analysis of telomeres in peripheral blood cells from patients with bone marrow failure. Pediatric Blood & Cancer, 2009 53(3): p. 411–416.

61. Imam, T., et al., Leukocyte Telomere Length in HIV Infected Pregnant Women Treated With Antiretroviral Drugs During Pregnancy and Their Uninfected Infants. JAIDS Journal of Acquired Immune Deficiency Syndromes, 2012 60(5): p. 495–502.

62. Zanet, D. L., et al., Blood and Dried Blood Spot Telomere Length Measurement by qPCR: Assay Considerations. PLOS ONE, 2013 8(2): p. e57787.

63. Jodczyk, S., et al., Telomere length measurement on the Roche LightCycler 480 Platform. Genet Test Mol Biomarkers, 2015 19(2): p. 63–8.

64. Panero, J., et al., Absolute qPCR for measuring telomere length in bone marrow samples of plasma cell disorders. Molecular biotechnology, 2015 57(2): p. 155–159.

65. Hsieh, A., et al., Optimization of a relative telomere length assay by monochromatic multiplex real time quantitative PCR on the LightCycler 480: sources of variability and quality control considerations. The Journal of Molecular Diagnostics, 2016 18(3): p. 425–437.

66. Lynch, S. M., et al., Race, Ethnicity, Psychosocial Factors, and Telomere Length in a Multicenter Setting. PLOS ONE, 2016 11(1): p. e0146723.

67. Wand, T., et al., Telomere content measurement in human hematopoietic cells: Comparative analysis of qPCR and Flow FISH techniques. Cytometry A, 2016 89(10): p. 914–921.

68. Behrens, Y., et al., Comparison of different methods for telomere length measurement in whole blood and blood cell subsets: recommendations for telomere length measurement in hematological diseases. Genes Chromosomes and Cancer, 2017 56(9): p. 700–708.

69. Khincha, P., et al., Correlation of leukocyte telomere length measurement methods in patients with dyskeratosis congenita and in their unaffected relatives. International journal of molecular sciences, 2017 18(8): p. 1765.

70. Lee, M., et al., Comparative analysis of whole genome sequencing based telomere length measurement techniques. Methods, 2017 114: p. 4–15.

71. Tarik, M., et al., Validation of quantitative polymerase chain reaction with Southern blot method for telomere length analysis. Future Science OA, 2018 4(4): p. FSO282.

72. Wang, Y., et al., Telomere Length Calibration from qPCR Measurement: Limitations of Current Method. Cells, 2018 7(11): p. 183.

73. Ferreira, M., et al., Comparison of flow FISH and MM qPCR telomere length assessment techniques for the screening of telomeropathies. Annals of the New York Academy of Sciences, 2019: p. 1–11.

74. Ropio, J., et al., Reliable blood cancer cells’ telomere length evaluation by qPCR. Cancer Med, 2020 9(9): p. 3153–3162.

75. Ropio, J., et al., Reliable blood cancer cells’ telomere length evaluation by qPCR. Cancer Medicine, 2020

76. Verhulst, S., Improving comparability between qPCR based telomere studies. Molecular, Ecology Resources, 2020 20(1): p. 11–13.

77. Verhulst, S., et al., Commentary: The reliability of telomere length measurements. International Journal of Epidemiology, 2015 44(5): p. 1683–1686.

78. Benetos, A., et al., Response by Benetos et al to letter regarding article,“Short leukocyte telomere length precedes clinical expression of atherosclerosis: the blood and muscle model”. Circulation research, 2018 122(8): p. e73–e74.

79. Benetos, A., et al., Telomere length tracking in children and their parents: implications for adult onset diseases. Faseb j, 2019 33(12): p. 14248–14253.

80. Dlouha, D., et al., Comparison of the relative telomere length measured in leukocytes and eleven different human tissues. Physiol Res, 2014 63 Suppl 3: p. S343–50.

81. Dagnall, C., et al., Effect of pre analytic variables on the reproducibility of qPCR relative telomere length measurement. PLOS ONE, 2017 12(9): p. e0184098.

82. Brazma, A., et al., Minimum information about a microarray experiment (MIAME)—toward standards for microarray data. Nature genetics, 2001 29(4): p. 365–371.

83. Taylor, C., et al., The minimum information about a proteomics experiment (MIAPE). Nature biotechnology, 2007 25(8): p. 887–893.

84. Field, D., et al., The minimum information about a genome sequence (MIGS) specification. Nature biotechnology, 2008 26(5): p. 541–547.

85. Taylor, C., et al., Promoting coherent minimum reporting guidelines for biological and biomedical investigations: the MIBBI project. Nature biotechnology, 2008 26(8): p. 889–896.

86. Fiehn, O., et al., The metabolomics standards initiative (MSI). Metabolomics, 2007 3(3): p. 175–178.

87. Brazma, A., Minimum information about a microarray experiment (MIAME)–successes, failures, challenges. The Scientific World Journal, 2009 9.

88. Kimura, M., et al., Measurement of telomere length by the Southern blot analysis of terminal restriction fragment lengths. Nature protocols, 2010 5(9): p. 1596–1607.

