## Supplemental Tables and Methods for "Method comparison studies of telomere length measurement using qPCR approaches: a critical appraisal of the literature"

Supplemental Table 1. Telomere Research Network Reporting Guidelines.

| Category (score) | Reporting Item | Importance | MIQE Importance |
|---|---|---|---|
| Sample type, storage, DNA extraction and integrity (score out of 9 or 10) | Sample type | E | E |
| | Sample storage temperature | E | E |
| | Sample storage time before extraction | D | E |
| | Sample storage buffer | D | E |
| | DNA extraction method | E | E |
| | DNA storage conditions | E | N/A |
| | DNA freeze-thaw cycles | E | N/A |
| | Method of documenting DNA quality/integrity | D | E |
| | % of samples tested for DNA quality/integrity | D | N/A |
| | **For studies with repeated measures design report the above for all time points* | E | N/A |
| qPCR assay (score out of 13 or 15) | State type of PCR method | E | N/A |
| | PCR machine | D | E |
| | Source of master mix & reagents | E | E |
| | Final reaction volume | E | E |
| | Telomere primer sequences | E | E |
| | Telomere primer concentration | E | N/A |
| | Single copy gene name | D | N/A |
| | Single gene primer sequence | E | E |
| | Singe gene primer concentration | E | N/A |
| | Full PCR program description including temperature, times, and cycle numbers | E | E |
| | PCR efficiency of single copy gene and telomere primers | E | E |
| | Source of control samples | E | N/A |
| | Concentration of DNA standard | E | N/A |
| | *For aTL PCR measurement only: sequence of oligo standards* | E | E |
| | *For aTL PCR measurement only: concentration of oligo standards* | E | N/A |
| Data analysis (score out of 12 or 13) | Mean and standard deviation or median and range of telomere lengths | D | N/A |
| | Number of sample replicates | E | E |
| | Level of independence of the replicates (plate vs day vs extraction) | E | E |
| | Analytic method, considering replicate measurements, to determine final TL | E | N/A |

| | | | |
|---|---|---|---|
| | Method of accounting for variation between sample replicates | E | N/A |
| | Method of accounting for well position effects | E | N/A |
| | Method of accounting for between-plate effects | E | N/A |
| | % of samples repeated due to failed QC | D | E |
| | % of samples excluded from analysis due to failed QC | E | E |
| | Acceptable range of PCR efficiency for primers | D | N/A |
| | ICCs of sample/study groups | E | N/A |
| | T/S ratio transformed to Z score prior to analysis | D | N/A |
| | *For studies with family samples or repeated measures design: analytic method to account for this* | E | N/A |

Supplemental Table 2. Morinha Reporting Guidelines.

| Category (score) | Reporting Item | Significance | MIQE Importance |
|---|---|---|---|
| Sample (score out of 5) | Experimental and control groups characteristics | A | E |
| | Tissue sampled | A | E |
| | Volume/mass of sample processed | B | D |
| | Storage conditions in the field and lab (including freeze-thaw cycles) | A | E |
| | Storage time before DNA extraction | B | E |
| DNA extraction (score out of 7) | Name of kit and details of any modifications | A | E |
| | Procedure and/or instrumentation | A | E |
| | Extraction method of the calibrator sample (if applicable) | A | N/A |
| | DNA quantification method | B | E |
| | DNA quality and purity (integrity, yield, 260/280 and 260/230 ratios) | A | D |
| | Storage conditions | A | N/A |
| | Freeze-thaw cycles before qPCR | B | N/A |
| qPCR validation (score out of 7) | Calibration curves with slope and y-intercept | A | E |
| | R2 of calibration curve | A | E |
| | qPCR efficiencies (telomere and reference genes) | A | E |
| | Cq of the NTC | B | E |
| | Linear dynamic range | B | E |
| | Specificity (e.g. gel, sequence, melt) | A | E |
| | Amplification and melting curve plots | B | N/A |
| qPCR protocol (score out of 12 or 13) | Primer sequences | A | E |
| | PCR Master mix (manufacturer and name of the products used in the assay) | A | E |
| | Complete reaction conditions (dNTP, MG2+, primer and polymerase concentrations, DNA amount, other components, reaction volume | A | E |
| | Additives (SYBR Green I, ROX, DMSO, etc) | A | E |
| | Calibrator sample used (e.g. synthetic, pool) | A | N/A |
| | Singleplex or multiplex | A | N/A |
| | *If singleplex, were both reactions run on the same plate or not* | A | N/A |
| | Number of replicates (technical and biological) | A | E |
| | Were different groups (age/sex/etc) run on the same plates or randomized | A | N/A |
| | Manufacturer of plates/tubes and catalog number | C | D |
| | Complete thermocycling parameters | A | E |
| | Reaction setup (manual/robotic) | C | D |
| | qPCR instrument | A | E |

| | | | |
|---|---|---|---|
| Data analysis (score out of 13) | Quality control steps for data | A | E |
| | Analysis program (source, version) | A | E |
| | Method used in the data analysis | A | E |
| | Choice of reference genes | B | E |
| | Concentrations of the calibrator sample used to create standard curves | A | N/A |
| | Normalalization and adjustment methods | A | E |
| | Repeatability (intra-assay variation) | A | E |
| | Reproducibility (inter-assay variation statistics) | A | D |
| | Acceptance and rejection criteria | A | E |
| | How was telomere length calculated? | A | N/A |
| | Statistical methods for results significance | A | E |
| | Software used for statistical analysis (source, version) | A | E |
| | Cq or raw data submission | B | D |

Supplemental Table 3. Reporting guidelines rubric.

| | |
|---|---|
| **For all reporting guidelines:** | ▪ Record information as reported only if it is provided directly in the paper itself or in the 1st order of references cited (with exceptions related to Cawthon papers – see below). Do not include information that extends beyond one previous citation. For example, if the methods section states that the detailed methods are described in a previous paper, report only the information from that cited paper. If the referenced paper itself has an additional reference for methods, mark this information as not provided.<br> o Applies for both methods and sample demographics (i.e. if further analysis is done on samples from a previously published study)<br>▪ Methods sections that state "followed Cawthon 2002 and/or 2009" and failing to provide any additional information about the specific assay performed are not counted as reporting assay parameters and conditions<br>▪ If sample demographics for the overall population in the study are provided, and only a subset of samples were involved in cross method analyses, reviewer can consider the demographic characteristics as present for the cross-method sample<br>▪ Information in supplemental material or supplemental methods is reported as present.<br>▪ qPCR efficiencies in qPCR assay/validation section: this refers to actual efficiencies of the assays performed |
| **TRN reporting guidelines:** | ▪ Studies without repeated measures design: do not include "for studies with repeated measures…." in score for sample type, storage, extraction, and integrity (mark N/A) or "method of accounting for within-family samples or repeated measures design" if it does not apply (mark N/A, exclude from score)<br>▪ Studies without aTL measurement: mark N/A for "for aTL PCR measurement…" and do not include in score<br>▪ DNA quality and integrity: report as yes if methods section describe how DNA quality was assessed; actual DNA values not required (note this differs from the Morinha requirement)<br>▪ Acceptable range of PCR efficiency: Note this differs from actual PCR efficiency values in the experiment, and should express the lab's typical acceptability/exclusion criteria for PCR assay efficiency |
| **Morinha reporting guidelines:** | ▪ Experimental and control group characteristics: if cross-method analysis is performed on only a subset of samples, report this as present only if characteristics are provided specifically for the subset of samples (not just the entire experiment)<br>▪ DNA quality and purity: report yes only if actual DNA integrity, yield, 260/280, 260/230 ratios are provided (ranges/mean are acceptable)<br>▪ Choice of reference genes: Name of single-copy gene |

Supplemental Table 4. Individual paper results of reporting guidelines.

| Author | TRN Reporting Guidelines | | | | Morinha Reporting Guidelines | | | | | | ***Average Overall Grade*** |
|---|---|---|---|---|---|---|---|---|---|---|---|
| | Sample | qPCR assay | Data analysis | ***Overall Grade*** | Sample | DNA extraction | qPCR validation | qPCR protocol | Data analysis | ***Overall grade*** | |
| Hsieh | 44% | 92% | 58% | 68% | 40% | 71% | 71% | 85% | 92% | 78% | **73%** |
| Eisenberg | 67% | 85% | 58% | 71% | 40% | 86% | 29% | 83% | 85% | 70% | **71%** |
| Zanet | 56% | 92% | 50% | 68% | 80% | 29% | 43% | 77% | 92% | 69% | **68%** |
| Martin-Ruiz (b) | 50% | 92% | 77% | 75% | 40% | 57% | 0% | 62% | 100% | 60% | **68%** |
| Tarik | 56% | 92% | 58% | 71% | 40% | 29% | 71% | 75% | 77% | 64% | **67%** |
| Jodcyzk | 67% | 92% | 58% | 74% | 40% | 43% | 0% | 69% | 77% | 53% | **63%** |
| Gutierrez-Rodrigues | 78% | 77% | 42% | 65% | 60% | 57% | 0% | 77% | 77% | 60% | **62%** |
| Ropio | 67% | 67% | 33% | 56% | 80% | 71% | 0% | 69% | 85% | 64% | **60%** |
| Imam | 33% | 77% | 33% | 50% | 80% | 29% | 14% | 77% | 69% | 58% | **54%** |
| Ehrlenbach | 30% | 46% | 69% | 53% | 20% | 43% | 0% | 62% | 83% | 49% | **51%** |
| Aviv | 33% | 85% | 25% | 50% | 40% | 14% | 0% | 69% | 69% | 47% | **48%** |
| Gadalla | 56% | 69% | 25% | 50% | 60% | 29% | 0% | 54% | 54% | 42% | **46%** |
| Khincha | 22% | 69% | 42% | 47% | 40% | 14% | 0% | 54% | 69% | 42% | **45%** |
| Wang | 22% | 69% | 42% | 47% | 40% | 14% | 0% | 46% | 54% | 36% | **41%** |
| Panero | 11% | 73% | 18% | 40% | 40% | 0% | 43% | 77% | 31% | 42% | **41%** |
| Behrens | 33% | 62% | 33% | 44% | 40% | 29% | 0% | 38% | 62% | 38% | **41%** |
| Lee | 11% | 69% | 17% | 35% | 40% | 0% | 0% | 69% | 38% | 36% | **35%** |
| Pavesi | 11% | 77% | 8% | 35% | 40% | 0% | 0% | 54% | 46% | 33% | **34%** |
| Wand | 11% | 46% | 25% | 29% | 40% | 29% | 0% | 38% | 54% | 36% | **32%** |
| Ventura Ferreira | 22% | 31% | 25% | 26% | 40% | 29% | 0% | 25% | 46% | 29% | **28%** |
| Gardner | 44% | 92% | 25% | 56% | 40% | 43% | 0% | 77% | 77% | 56% | **56%** |
| Hunt | 11% | 92% | 33% | 50% | 40% | 0% | 0% | 85% | 85% | 53% | **52%** |
| Martin-Ruiz (a) | 22% | 85% | 33% | 50% | 20% | 33% | 0% | 69% | 31% | 36% | **43%** |
| Salpea | 22% | 92% | 50% | 56% | 40% | 29% | 71% | 69% | 85% | 60% | **58%** |
| Lynch | 33% | 46% | 42% | 41% | 80% | 29% | 0% | 78% | 69% | 49% | **45%** |
| **Average per Category** | **37%** | **75%** | **39%** | **52%** | **46%** | **32%** | **14%** | **66%** | **68%** | **50%** | **51%** |

Supplemental Methods. Calculation of the repeatability (ICC) of telomere length measures.

Unlike the CV statistics, the ICC cannot be calculated for each biological sample individually, but instead is calculated for a set of biological samples. Thus, to be able to calculate the repeatability, (a subset of) samples should be re-measured one or more times in an identical fashion as TL is typically measured in one's laboratory. Two important considerations when selecting these samples are the following. Firstly, variation in TL measurement can arise at any point between sample collection and arrival at your TL estimate. Therefore, the ICC will be overestimated when, for example, the ICC is estimated over repeated measurements of extracted DNA relative to measurements where the DNA extraction is also repeated (but by how much is not known). Thus, the ICC you estimate from the data will be closer to the true ICC when more of the measurement process is repeated independently in the successive measurements. Secondly, the TL of the subset of samples needs to be representative of the complete set of samples that is analysed. This is true in particular for the range of TL in the sample – having a larger range in the subset than in the ultimate sample will overestimate the 'true' ICC, while having a smaller range in the subset than in the ultimate sample will result in underestimation of the ICC.

The ICC can be calculated in different ways, depending on how variation between batches is accounted for (i.e. not at all, as random effect, or as fixed effect), where 'batches' can be thought of as different plates, or gels, or measurement sessions that differ in any respect (e.g. day, person, equipment, laboratory). The best way to account for batch in the ICC calculation is to use the same approach as in the ultimate analyses in which hypotheses are tested.

The text below is an R script, mixing instructions with the actual script. When a line is precede by '#', this indicates it is a comment – and it will not be executed. It is left in the text here to make it possible to copy all text below (up to the References) to an R-script – see instructions below.

```
# R script to calculate the ICC (IntraClass Correlation), also known as 'repeatability'.
# The text below assumes you are new to R....
# '#' before text indicates it is a comment - will not be executed
# The other lines need to be 'run' and results will show in the console window.

# Before you start
# 1. it is advisable to do the analysis in Rstudio (freely downloadable and works on
# multiple platforms), which serves as a 'shell' to R, and copy this text to a new R-script.
# 2. A useful introduction by the authors of the package used in this script to calculate
# the ICC is recommended reading:
# https://cran.r-project.org/web/packages/rptR/vignettes/rptR.html
# The text below is only enough to get you started
# 3. Data format
# The data need to be in the 'long' format. This implies ALL the telomere estimates are
# in one column, with sample identity in another column. Data may have been collected in
# different 'batches' (plates, gels, days, labs) and batch identity is coded in a separate
# column.
# When your data is in wide format, this can easily be changed to a long format in Excel.
# So the data file usually has a minimum of three columns (with variable names in brackets):
```

```
# sample identity (id), telomere estimate (TL), batch (batch) [when you use different
# variable names, the names in the code below need to be replaced with those names].

# The packages listed below are needed and you will probably need to install them first.
# At the top of the bottom right panel in RStudio there is a tab 'packages' you can use.
library(readxl) #for when your data are in Excel format
library(MASS)
library(lme4)
library(rptR)

# To clear lists of objects – useful to run whenever you start an analysis:
rm(list=ls())
rm(list = ls(all = TRUE))

# Reading in the data.
# In the example below, the data were stored in Excel, but Rstudio reads many formats.
# Other formats may require loading another package - Rstudio will tell you this.

# Note that the first bit of the code below, 'd <-', you can read as 'd becomes'.
# We here arbitrarily name the data set 'd' (commands / names in R are case sensitive!).

# You can get the import code for your file location and correct command for your file format
# using the 'import dataset' tab in RStudio (above the top right frame on a mac).

# When you import a data set using RStudio, the dataset will have a name different from
# what is in the script below. I recommend copying the code you see after the "<-" to the
# script just before actually importing the data and copy this below after the text "d <-".
# Alternatively, you replace 'd' in the code with the name you have given your data set.

d <- read_excel("FILE LOCATION/example.xls") #importing the example data from Excel file
# The file 'example.xls' is available on the telomere network site.

# Calculating the ICC
# The ICC can be calculated in different ways, depending on how variation between
# batches is accounted for (i.e. not at all, as random effect, or as fixed effect).
# The best way to account for batch in the ICC calculation is to use the same approach
# as in the analysis for which the data were collected.
# Below is the code for different ways to include 'batch' in the analysis.

# When running the scripts below, depending on details,
# there may be 'Singularity' issues that are reported as errors.
# You can safely ignore these (see information on rptR package for details.)

# 1. No correction for batch
```

```
# Including 'id' only - i.e. batch is not in the model
# Note that the (1|something) codes for a random intercept for levels of 'something'
rpt(TL ~ (1|id), grname = "id", data=d, datatype = "Gaussian", nboot = 1000, npermut=0)

# 2. Including 'batch' as random effect
rpt(TL ~ (1|id) + (1|batch), grname = "id", data=d, datatype = "Gaussian", nboot = 1000, npermut=0)

# 3. Including 'batch' as fixed effect
rpt(TL ~ batch + (1|id), grname = "id", data=d, datatype = "Gaussian", nboot = 1000, npermut=0)

# Note that these models can be extended with other factors and covariates.
# See example below that includes age (as fixed effect)
rpt(TL ~ age + (1|id) + (1|batch), grname = "id", data=d, datatype = "Gaussian", nboot = 1000,
npermut=0)
# When running the model including age, the ICC is likely to become lower. The new (lower)
# ICC estimate is the more relevant estimate. This is so, because you will probably
# take age into account in your statistical analyses also. The remaining variation in the
# data will decrease as a consequence, and the ICC is calculated over the remaining variation.
# Adding other factors to the model, e.g. batch identity or procedure characteristics
# that you also include in the model with which you test hypotheses can also be added as
# factors to the model, and may increase the ICC.

# The script above assumes the TL measurements are normally distributed, but the
# rptR package can handle other distributions.

# Extrapolated repeatability – an important extension!
# Suppose your protocol includes rTL measurement in duplicate, using for example two plates
# with samples in triplicate on each plate, and you use the average of the two plates in
# the data analysis. You can then calculate the ICC over the two plates, but this will
# underestimate the ICC of the average of the two plates, which is based on more measurements.
# However, the ICC of the mean of the two plates can be calculated with a simple equation
# once the ICC over the two plates is known:  when r is your repeatability (ICC) estimate,
# and n is the number of replicate measurements then the extrapolated repeatability (re):
# re = r / (r+1/n*(1-r))
# This is equation 37 in: Nakagawa, S. & Schielzeth 2010, Biological Reviews 85, 935-956.
```

Supplemental Information Legends.

**S1 Table. Telomere Research Network (TRN) Reporting Guidelines.** Reporting items of the TRN Guidelines by category, with assigned importance for each item and comparison with the related assigned importance of the MIQE guidelines. E: essential information, should be submitted with the manuscript. D: desired information, should be submitted if available.

**S2 Table. Morinha Reporting Guidelines.** Reporting items of the guidelines developed by Morinha et al. by category, with assigned importance for each item and comparison with the related assigned importance of the MIQE guidelines. A: very important. B: somewhat important. C: somewhat important, should be submitted if available. E: essential information, should be submitted with the manuscript. D: desired information, should be submitted if available.

**S3 Table. Grading rubric for TRN and Morinha Reporting Guidelines.**

**S4 Table. Results of reporting guidelines for individual papers.** Scores for each category are provided as a percentage of items in that category sufficiently reported. A total score for each set of reporting guidelines are calculated from all items as well as an average overall score between the two reporting guidelines.

**Supplemental Methods. Calculation of the repeatability (ICC) of telomere length measures**
